# Antibiotic dose-response curves can measure antibiotic activity against *Mycobacterium abscessus* and *Mycobacterium peregrinum*

**DOI:** 10.64898/2025.12.16.694619

**Authors:** Husain Poonawala, Kathleen Davis, Myles E. Kenny, Ares Alivisatos, Nhi Van, Tracy Washington, Vinicius Calado Nogueira de Moura, Charles L. Daley, Bree B. Aldridge

**Author notes:** Now at Tufts University School of Medicine, Boston, MA. Now at UMass Chan Medical School, Worcester, MA. Corresponding author: Husain Poonawala.

## Abstract

*Mycobacterium abscessus* is a drug-resistant pathogen associated with poor clinical outcomes despite treatment with multidrug antibiotic regimens. Apart from clarithromycin, antimicrobial susceptibility testing (AST) results for *Mycobacterium abscessus* cannot guide antibiotic selection. AST involves measuring the minimum inhibitory concentration (MIC), which is allowed to span a four-fold range in concentration. This accepted variability of the MIC limits the clinical utility of AST. Antimicrobial dose-response curves, obtained by measuring the growth inhibition of a given organism to increasing concentrations of an antibiotic, can yield metrics of antibiotic activity that are less variable than the MIC. We used Clinical and Laboratory Standards Institute growth conditions for rapid-grower nontuberculous mycobacteria to generate 990 dose-response curves across three time points (72 hours, 96 hours, and 120 hours) for six guideline-recommended (clarithromycin, amikacin, cefoxitin, linezolid, tigecycline, and clofazimine) and five new (omadacycline, tedizolid, SPR719, SQ109, and bedaquiline) antibiotics against *Mycobacterium abscessus* subspecies *abscessus* ATCC 19977 and *Mycobacterium peregrinum* ATCC 700686. We established the fit of the dose-response curve (R^2^) as a quality control metric. Using the geometric standard deviation and median coefficient of variation, we demonstrated that the IC_50_ and IC_75_ (antibiotic concentrations corresponding to 50% and 75% growth inhibition, respectively) are less variable than the MIC. We identified time-dependent changes in dose-response curve metrics that allow the detection of inducible clarithromycin resistance with only five days of incubation. This study demonstrates the potential of dose-response curves in measuring antibiotic activity against *Mycobacterium abscessus*.

## Introduction

*Mycobacterium abscessus* (MAB) is a highly drug-resistant nontuberculous mycobacteria (NTM) responsible for difficult-to-treat pulmonary and extrapulmonary infections.^1,2^ The 2020 multi-society guidelines for the treatment of NTM pulmonary disease recommend treatment with a combination of two to six drugs, with the caveat that “the optimal drugs, regimens, and duration of therapy are not known.”^1,3^ Most patients will experience side effects from one or more antibiotics, often resulting in treatment cessation; fewer than half will attain microbiological cure.^4–11^

Treatment regimens for MAB are designed using the results of antimicrobial susceptibility testing (AST) performed according to Clinical and Laboratory Standards Institute (CLSI) guidelines for rapid-grower NTMs.^3,12^ However, with the exception of clarithromycin, MAB AST results are not predictive of clinical outcomes.^8,12^ Most MAB isolates demonstrate inducible clarithromycin resistance, which is associated with treatment failure, and is identified using a combination of methods available only in reference laboratories.^1,13^ Of the remaining six drugs recommended for the treatment of MAB, AST results are either not predictive of treatment outcomes (amikacin, cefoxitin, and linezolid), unreliable due to instability (imipenem), or without interpretive criteria (clofazimine and tigecycline).^1,14–17^ There are no breakpoints for omadacycline, tedizolid, or bedaquiline, newer drugs that are promising for the treatment of MAB infections.^1,18,19^ The complexity and inadequacy of AST contribute to the lack of progress against MAB.^1^

AST involves measuring the minimum inhibitory concentration (MIC) — the lowest antibiotic concentration required to limit visible growth of an organism — and is specific to any antibiotic-bacterial pair.^20^ For NTMs, AST is performed using broth microdilution.^12^ MICs are measured with 2-fold increases in concentration, and because measuring them involves numerous sources of error, repeat measurements can be one 2-fold measurement above or below the original measurement.^21^ This inherent variability of the MIC limits its use in antibiotic dosing, as seen with vancomycin (area under the curve (AUC) over 24 hours/MIC) for the treatment of methicillin-resistant *Staphylococcus aureus*.^22–24^ The variability in measuring the MIC has also made it challenging to use AST results to predict clinical outcomes, especially in patients with immunocompromising conditions, those receiving multiple antibiotics, or oral antibiotics – all of which apply to MAB infections.^21^ New metrics of antibiotic activity that are precise and predictive of clinical outcomes are needed.

An antibiotic dose-response curve can be generated by measuring microbial growth response to increasing concentrations of an antibiotic.^25,26^ This allows for a richer estimation of *in vitro* antibiotic efficacy using metrics such as the IC_50_ (antibiotic concentration at 50% growth inhibition) and the Hill slope, which are not only more precise than the MIC, but also have the potential to individualize antibiotic dosing.^22,26^ Reliable *in vitro* results might predict antibiotic-specific clinical outcomes, thereby allowing the development of more effective and less toxic treatments for drug-resistant infections and ultimately leading to improved patient outcomes.^22^

In this paper, we describe the use of CLSI rapid-grower NTM AST conditions to generate dose-response curves for six first-line and five novel antibiotics for MAB.^12,19^ We demonstrate that antibiotic dose-response curves can be used to obtain reliable metrics for the *in vitro* activity of clinically important drugs and detect inducible resistance to clarithromycin in MAB at five days instead of fourteen.

## Methods

### Antibiotics

Antibiotics were obtained from MedChemExpress (Monmouth Junction, New Jersey) (bedaquiline, cefoxitin sodium, linezolid, SPR719, SQ109, tedizolid, tigecycline) and Thermo Scientific Chemicals (amikacin, clarithromycin, clofazimine). Omadacycline was provided by Paratek Pharmaceuticals (King of Prussia, Pennsylvania). Single-use antibiotic stock (concentrations in Table S1) aliquots of 10-20 μL were prepared using dimethylsulfoxide (DMSO; bedaquiline, clarithromycin, linezolid, SPR719, SQ109, tigecycline, tedizolid) or sterile water with 0.01% Triton-X (amikacin, omadacycline) and stored at −70°C. Cefoxitin sodium (sterile water with 0.01% Triton-X) and clofazimine (DMSO) were prepared on the morning of each experiment. Antibiotics were allowed to come to room temperature prior to dispensing. Clofazimine was ultrasonicated for 10 minutes to facilitate dissolution.

### Strains and media

*Mycobacterium abscessus* subspecies *abscessus* ATCC 19977 (MAB) and *Mycobacterium peregrinum* ATCC 700686 (MPER) were used for all experiments. Frozen aliquots of isolates were subcultured onto 7H10 or 7H11 plates and incubated at 37***°*** C for 5-10 days until there were visible colonies. Similar-looking colonies were picked using a sterile loop and added to 100-200 μL of 7H9 Middlebrook media (prepared without Tween 80 or tyloxapol) in tubes pre-filled with 3mm zirconium beads (Ops Diagnostics, Lebanon, New Jersey) and vortexed for 20-30 seconds. These vortexed cells were then added to inkwells containing 5mL (MAB) or 3mL (MPER) of 7H9 media and incubated overnight in a shaking incubator (140 revolutions per minute) at 37***°*** C. Fresh cation-adjusted Mueller Hinton broth (CA-MHB) was prepared on the morning of each experimental run (as recommended by CLSI M07) by the Tufts Microbiology Media Kitchen.^27^

### Experimental procedure

Antibiotic stock solutions were dispensed into the 308 non-edge wells of clear, flat-bottomed 384-well microplates using a digital drug dispenser (D300e digital dispenser; Hewlett-Packard) as described previously.^28,29^ The 76 edge wells were filled with media only. For each antibiotic, the software was programmed to dispense 10 or 13 doses of specified volumes of antibiotic to obtain the desired antibiotic concentration in each well. Doses were spaced at 2-fold volume increments with the antibiotic concentration required to achieve 50% growth inhibition (IC_50_) at either the 5th or 6th dose (10-dose experiments) or 8th or 9th dose (13-dose experiment). Two replicates for each antibiotic were dispensed to determine technical variability. Drug wells were randomized on each plate to minimize confounding due to plate effects.

We adapted the rapid-grower NTM AST procedure for our procedure.^17^ For each isolate, 1.5 mL of an overnight culture was transferred from the inkwells into a microcentrifuge tube with zirconium beads and vortexed for 20-30 seconds. The optical density at 600 nm wavelength (OD_600_) was measured, and the culture was diluted to prepare a 0.2 OD_600_ suspension (equivalent to 0.5 McFarland Standard; confirmed by colony count) in 500 to 1000 µL of CA-MHB. The final inoculum was created by adding 100 µL of 0.2 OD_600_ suspension to 20mL of CA-MHB and 50 µL of this inoculum into each of the 308 non-edge wells.

Plates were sealed with an optically clear plate seal (AB-1170; Thermo Fisher Scientific), and OD_600_ was measured using a microplate reader (BioTek). To mix the antibiotic and inoculum, plates were centrifuged for 10 seconds at 480g in a plate centrifuge (FastGene Plate Centrifuge, Nippon Genetics Europe GmbH, Duren, Germany). They were then placed in a sealed plastic bag stacked with a maximum height of 3 plates and incubated at 30°C for 5 days.

### Plate reads and generation of dose-response curves

OD_600_ was measured for each plate on days 3, 4, and 5 using the optical plate reader. Plates were spun in the plate centrifuge (20-60 seconds at 480g), and seals were changed before each plate read to eliminate condensation under the seal that could interfere with OD_600_ measurement. The plate reads were processed, and dose-response curves were generated using custom scripts, as detailed previously.^28,29^ Printer layouts were used to derandomize plate reads to allow organization of OD_600_ reads by antibiotic. The median OD_600_ of the edge wells (OD_bkgd;_ containing only media) was subtracted from the raw OD_600_ values of non-edge wells to calculate the background-adjusted OD_600_ for all wells. The growth inhibition (G*_i_*) at any given antibiotic concentration was calculated by normalizing the bacterial growth at the corresponding concentration (OD_trt_) and normalizing it to the median OD_600_ of the untreated wells (OD_unt_), and subtracting it from 1, as shown in the equation below.

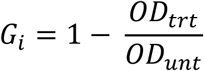

The resulting range of growth inhibition values corresponding to increasing concentrations of antibiotic was used to fit dose-response curves with a three-parameter Hill function using either the Levenberg-Marquadt algorithm or the trust-region-reflective algorithm; the results from the fit with the higher R^2^ value were chosen.^28^ The quality of each curve fit was assessed using a combination of the R^2^ and visual inspection of the curve. The MIC was recorded as the lowest concentration at which 100% growth inhibition was measured. The dose-response curve was used to obtain antibiotic inhibitory concentrations (IC) at 10% (IC_10_), 25% (IC_25_), 50% (IC_50_), 75% (IC_75_), 90% (IC_90_), and 95% (IC_95_) growth inhibition, respectively.^30^ The Hill function was used to obtain the area under the dose-response curve at 25% growth inhibition (AUC_25_), the Hill slope, and the maximum effect achievable (E_inf_).

### Analysis of metrics

We measured the variability of the MIC and IC values using two different methods. The first method used a range spanning the twofold concentration above and below the geometric mean of the MIC, as recommended by the CLSI.^20^ The second method used a range that was one geometric standard deviation above and below the geometric mean. To compare the variability between different IC measurements within and across different antibiotic-bacterial pairs, we calculated the median coefficient of variation using the ratio of the median absolute deviation (for any given metric) to the corresponding median.^31^ The accuracy of the MIC measurements was compared to CLSI QC ranges (MPER) or breakpoints (MAB).^12^ For antibiotics not covered under CLSI guidelines (bedaquiline, omadacycline, tedizolid, tigecycline, and SPR719) we used values from studies that used CLSI rapid-grower AST methodology; there were no values available for clofazimine or SQ109.^32–35^ We also calculated the median for the AUC_25_, E_inf_, and Hill slope.

Analyses were performed in R (version 4.5.2) and Rstudio (version 2025.9.1.401) using the tidyverse (version 2.0.0)suite of packages^36–38^

## Results

We performed experiments using eight biological replicates for each species (MPER and MAB). For each biological replicate, we obtained measurements from two technical replicates for all experimental runs (except one) with 15 replicates per antibiotic, except for clofazimine (13 replicates) and SQ109 (16 replicates). We generated 990 dose-response curves across these 66 combinations of species, antibiotics, and time points. An example of a dose-response curve and the metrics that can be obtained from it are shown in Fig. S1. The MICs were calculated for all antibiotic-bacterial pairs with the data used to generate dose-response curves and are discussed in the supplementary data and figures (Table S2, Fig. S1-S3)

### The R^2^ is a reliable measure of dose-response curve quality

The R^2^ (ranging between 0 and 1) compares the fitted dose-response curve (using three-parameter Hill curves) with the observed growth inhibition and was used to measure the quality of the fit. The median R^2^ value was greater than 0.9 for 58 of 66 antibiotic-bacterial-time-point combinations (Fig. 1A and 1B, and Table S2). The lowest median R^2^ values observed were for MPER-bedaquiline (all three time points) and MAB-bedaquiline and MAB-SQ109 (96 hours and 120 hours).

**Figure 1:**
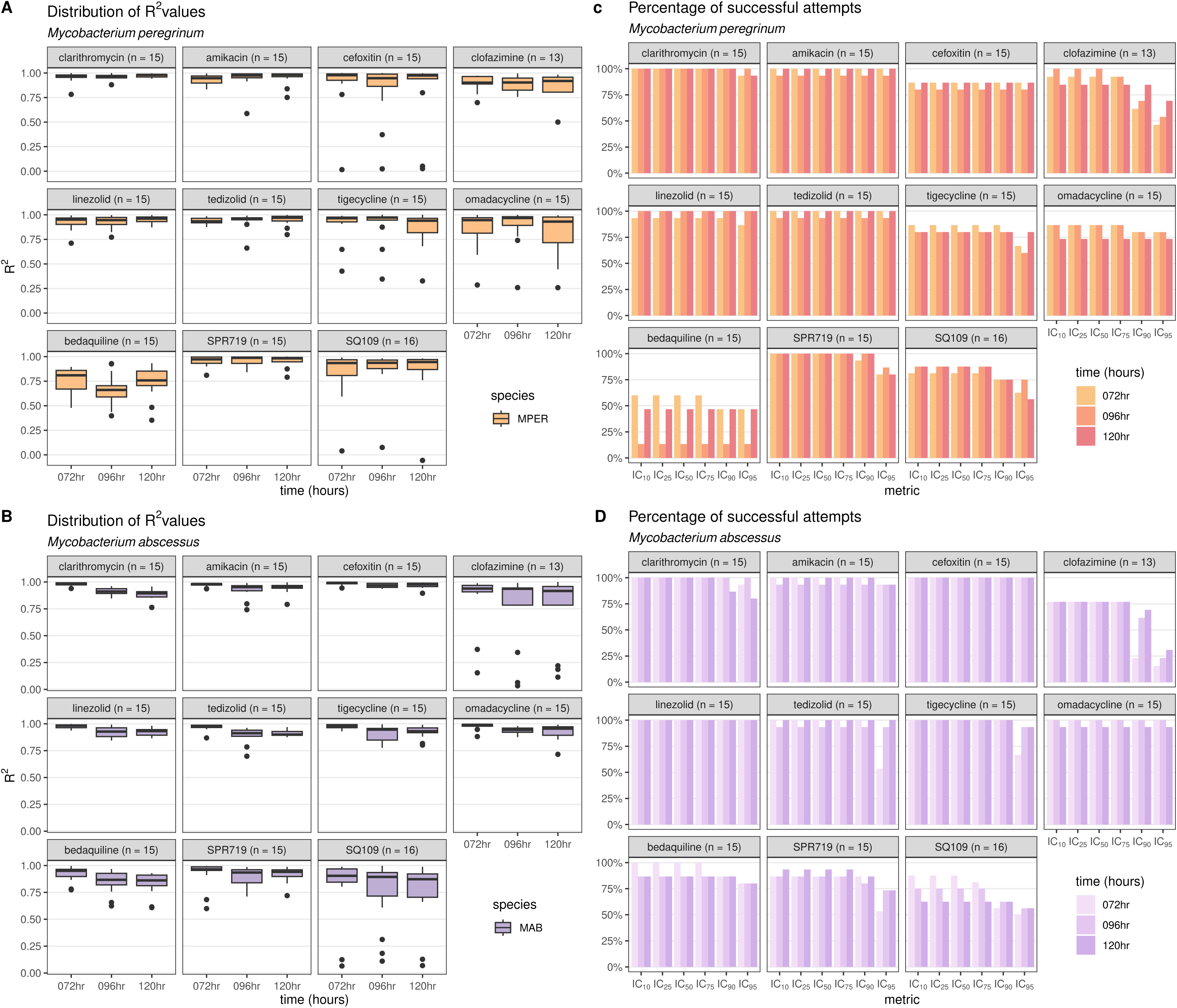
Assessment of dose-response quality. R^2^ values measure the fit of the dose-response curve with the observed growth inhibition and range from 0 (no agreement) to 1 (complete agreement). Boxplots of the R^2^ of each replicate (number of attempts in parentheses) for each antibiotic for *Mycobacterium peregrinum* (A) and *Mycobacterium abscessus* (B). The percentage of inhibitory concentration (IC) metrics that could be obtained from the fitted dose-response curve for each antibiotic (number of attempts in parentheses) at each time point was calculated for *Mycobacterium peregrinum* (C) and *Mycobacterium abscessus* (D). MPER, *Mycobacterium peregrinum*; MAB, *Mycobacterium abscessus* (B); hr, hours; IC_10,_ inhibitory concentration at 10% growth inhibition; IC_25_, inhibitory concentration at 25% growth inhibition; IC_50_, inhibitory concentration at 50% growth inhibition; IC_75_, inhibitory concentration at 75% growth inhibition; IC_90,_ inhibitory concentration at 90% growth inhibition; IC_95_, inhibitory concentration at 95% growth inhibition.

Antibiotic-bacterial pairs with low median R^2^ values were likely to have a lower percentage of successful attempts at obtaining IC values (Fig. 1). For example, for MPER-bedaquiline, the median R^2^ values ranged from 0.66 to 0.81, with the corresponding percentage of acceptable dose-response curves ranging from 13% to 60%. In contrast, all MAB-cefoxitin replicates had acceptable dose-response curves, with median R^2^ values of 0.99, 0.97, and 0.98 at 72 hours, 96 hours, and 120 hours, respectively. For both MPER and MAB, there were fewer IC_90_ and IC_95_ measurements than IC_10_-IC_75_ measurements for clofazimine, tigecycline, SPR719, and SQ109 (Fig. 1C and Fig. 1D). Except for clofazimine (MPER and MAB), bedaquiline (MPER), and SQ109 (MAB), the percentage of successful measurements remained consistent at each time point.

These data suggested that the R^2^ could be used as a proxy for quality, with dose-response curves with low R^2^ values being unlikely to be of acceptable quality or yield IC values. Filtering out dose-response curves with R^2^ values below 0.75 and that did not pass visual inspection improved the median R^2^ to 0.85 for all conditions (Table S2). After this filtering step, 888 dose response curves (89.7%) were available for further analysis.

### IC values are less variable than the MIC

We determined the variability of IC metrics using either a) ±1 2-fold concentration or b) 1 geometric standard deviation around the geometric mean of all replicates for any given metric (Table S2). We posited that the more precise the IC value or MIC, the narrower the spread of the geometric standard deviation around the geometric mean. Using cefoxitin as an example (Fig. 2), only 1-2 values were outside the 2-fold range (limits shown with red triangles) of the geometric mean (horizontal black line) for IC_10_, IC_25_, and IC_95_ for MPER and MAB. For all IC values, the range of the geometric mean and 1 geometric standard deviation (blue rectangle) was less than the 2-fold doubling range around the corresponding geometric mean (Fig. 2). In contrast, for the MIC, the range of the geometric mean and geometric standard deviation was greater than the 4-fold range of the geometric mean for both MAB and MPER.

**Figure 2:**
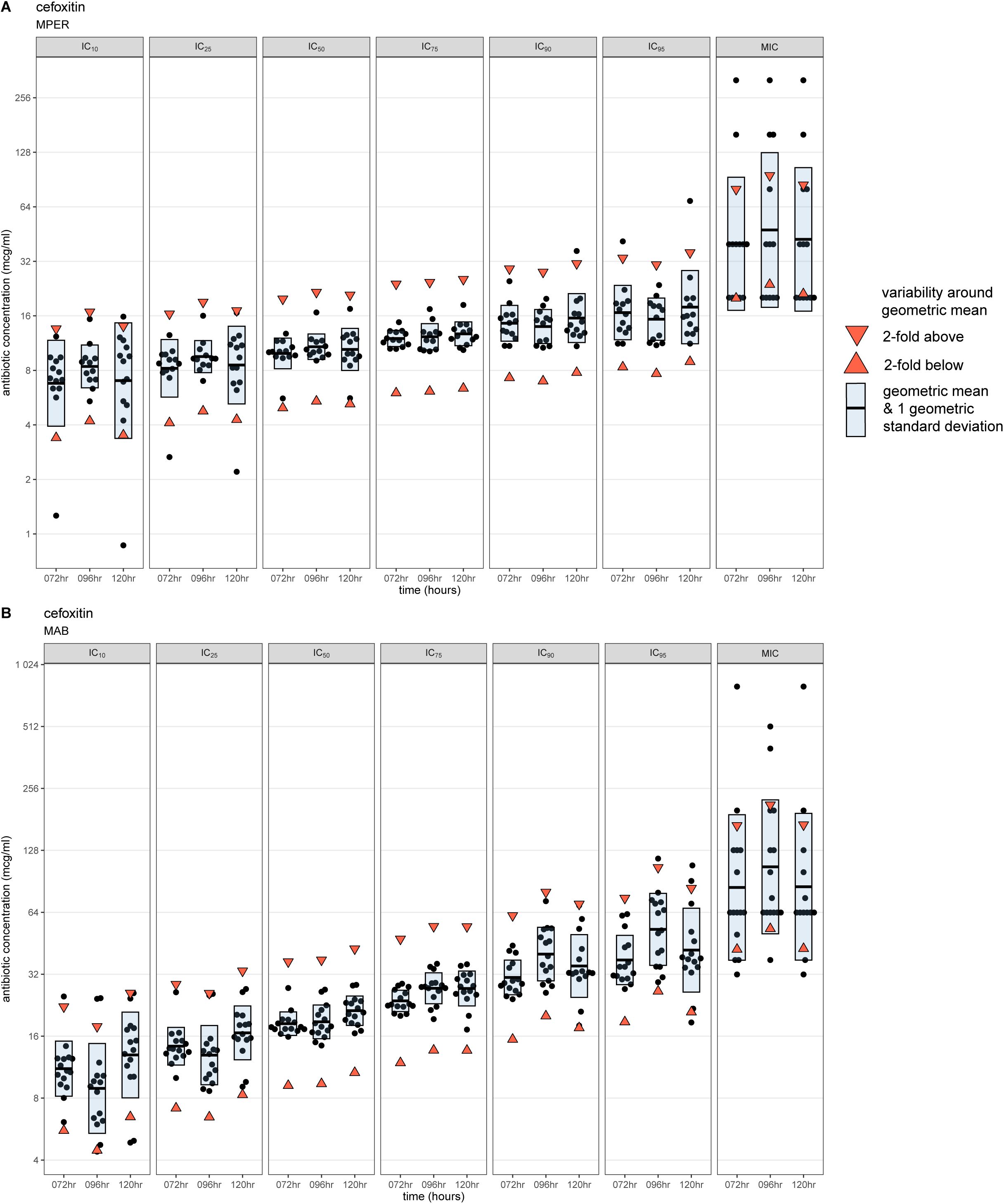
Variability of IC metrics and MIC for cefoxitin. The inhibitory concentration (IC) values for each replicate and the minimum inhibitory concentrations (MICs) for cefoxitin at each time point (black points) for *Mycobacterium peregrinum* (A) and *Mycobacterium abscessus* (B), with the geometric mean represented by the horizontal bar. The upper and lower limits of ±1 2-fold range around the geometric mean are represented by the downward- and upward-pointing red triangles, respectively. The blue shaded area represents 1 geometric standard deviation around the geometric mean. MPER, *Mycobacterium peregrinum*; MAB, *Mycobacterium abscessus* (B); hr, hours; IC_10,_ inhibitory concentration at 10% growth inhibition; IC_25_, inhibitory concentration at 25% growth inhibition; IC_50_, inhibitory concentration at 50% growth inhibition; IC_75_, inhibitory concentration at 75% growth inhibition; IC_90,_ inhibitory concentration at 90% growth inhibition; IC_95_, inhibitory concentration at 95% growth inhibition; MIC, minimum inhibitory concentration.

The reproducibility of IC metrics varied across antibiotic-bacterial pairs (Figs. 2 and S4-S6). Except for MPER-bedaquiline, MAB-tigecycline, MAB-SPR719, and MAB-SQ109, most replicates for IC_50_ and IC_75_ were within ± 2-fold concentration (range between red triangles) of the geometric mean (horizontal black line) and the spread of 1 geometric standard deviation around the geometric mean (blue shaded area) was less than the 2-fold concentration range around the geometric mean (range between red triangles). Across antibiotic-bacterial pairs and time points, we noted greater variability around the geometric mean for the IC_10,_ IC_25_, IC_90_, and IC_95_ than for the IC_50_ and IC_75._

### The IC_50_ and IC_75_ values are the least variable IC values

Because the median IC and MIC values approximated the geometric mean (Table S2), we used the median coefficient of variation (the non-parametric equivalent of the coefficient of variation) to compare the variability across different IC values and different antibiotic-bacterial pairs. A median absolute deviation equal to the median IC would be equivalent to a ±1 2-fold concentration around the median IC and result in a median coefficient of variation of 1. We therefore defined a median coefficient of variation of 1 or lower as an acceptable limit of variability.

Except for MPER-bedaquiline, the median coefficient of variation was less than 1 for most IC values for all antibiotic-bacterial pairs; the IC_10_ was the IC value for which the median coefficient of variation most frequently exceeded 1 (Fig. 3). We found that the median coefficient of variation was lowest for the IC_50_ and IC_75_ than for other IC values at any given time point, demonstrating a U-shape (Fig. 3). This U-shaped pattern was best seen with cefoxitin and was present across most antibiotic-bacterial pairs (Fig 3). Exceptions to this U-shaped pattern included clofazimine and tigecycline (both MPER and MAB) and bedaquiline-MPER, for which there were both lower median R^2^ values and lower percentage of successful attempts at obtaining IC values (Fig. 1 and Table S2). Antibiotic-bacterial pairs with lower median or wider spread of R^2^ values tended to have higher median coefficients of variation across all IC values, while preserving the U-shaped pattern to varying degrees, as seen with SPR719 and SQ109 (Fig. 3).

**Figure 3:**
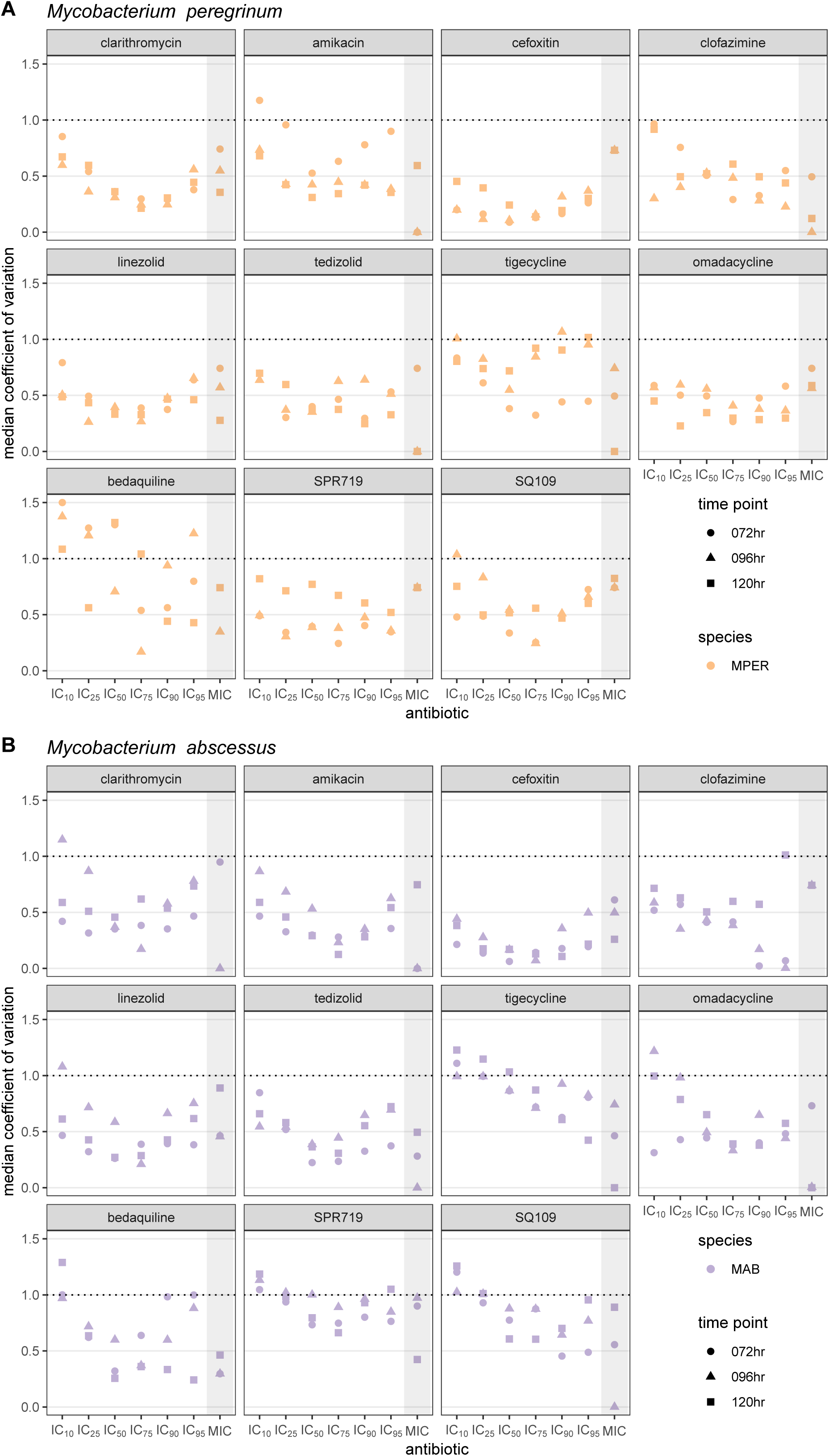
Comparison of the median coefficient of variation across IC and MIC values, time points and antibiotics. The median coefficient of variation allows comparison of variability across multiple dimensions and was calculated for *Mycobacterium peregrinum* (A) and *Mycobacterium abscessus* (B). We used a value of 1 or lower as an acceptable amount of variability as it represents the value at which the median standard deviation is equal to the median and would be equivalent to ±1 2-fold range around the median. MPER, *Mycobacterium peregrinum*; MAB, *Mycobacterium abscessus* (B); hr, hours.

### Dose-response metrics can identify inducible resistance to clarithromycin

The shape of MAB-clarithromycin (Fig. 4B) dose-response curves (dashed lines) changed over time; these changes were absent in MPER-clarithromycin (Fig. 4A). We obtained fewer MIC readings at 96 hours than at 72 hours, and none at 120 hours (Fig. 5B). On plotting the IC values, we noted the IC_50_ value remained unchanged across all time points and the IC_75_, IC_90_, and IC_95_ values increased with successive time points (Fig. 5B). Similarly, the AUC_25_, a measure of antibiotic potency, increased over time (Fig. 5C), consistent with increasing concentration of antibiotic necessary to achieve the same amount of growth inhibition at successive time points. On the other hand, the Hill slope decreased over time (Fig. 5E), suggesting that achieving complete growth inhibition would require a higher concentration of the antibiotic at later time points. All these metrics remained unchanged over time in MPER (Fig. 5A, C-E). We believe that these findings are likely explained by inducible resistance mediated by the *erm*(41) gene in the MAB ATCC type strain used in our experiments; this finding was not observed in MPER, which is not known to harbor any of the *erm* genes responsible for inducible resistance to clarithromycin.^17^

**Figure 4:**
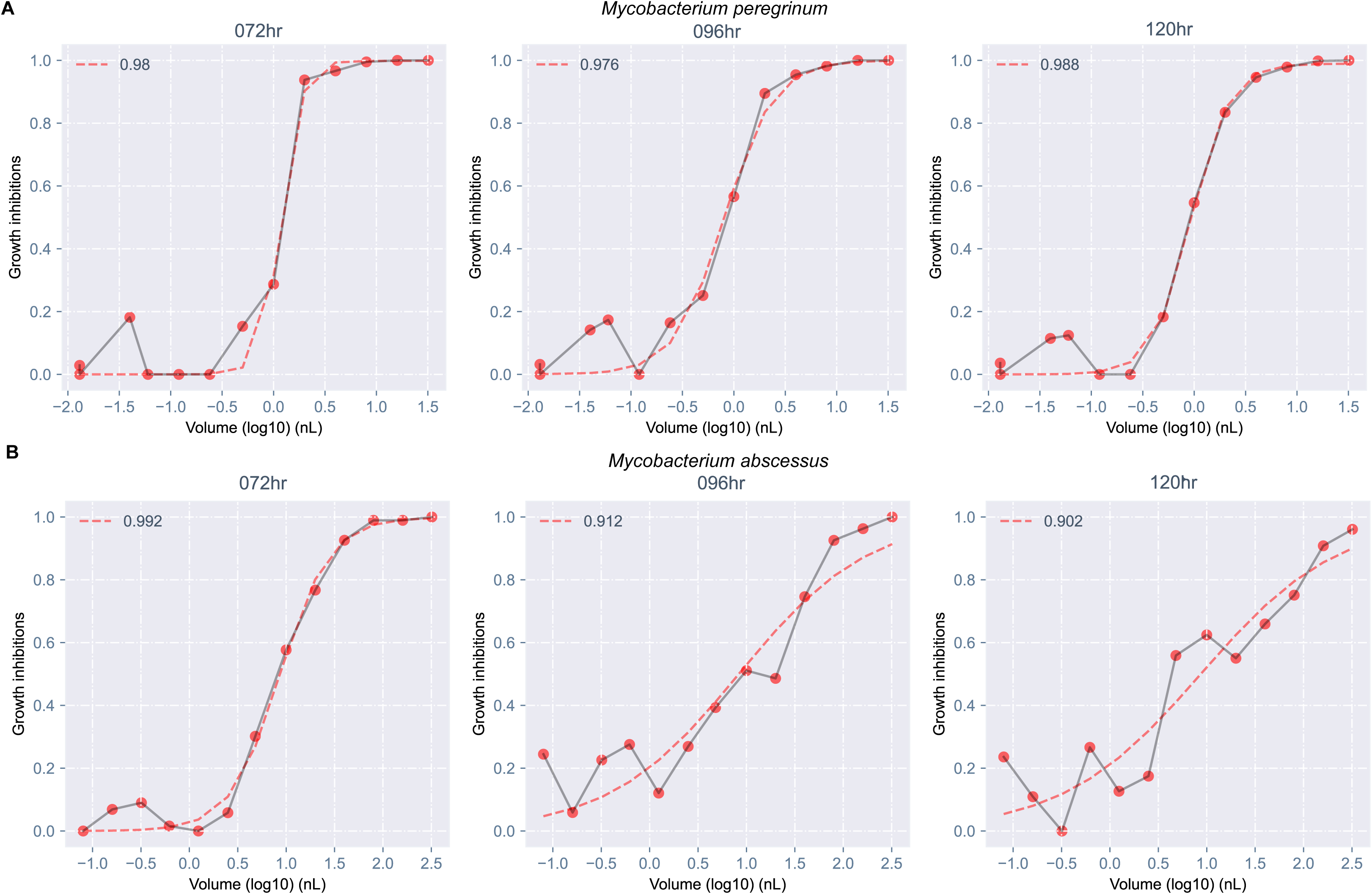
Temporal changes in dose-response curves consistent with inducible resistance to clarithromycin in *Mycobacterium abscessus*. Dose-response curves for clarithromycin showed no change in shape across time for *Mycobacterium peregrinum* (A). However, changes in shape were seen for *Mycobacterium abscessus* (B). The red points show calculated growth inhibitions at each concentration which are connected by a gray line. The red dashed line is the dose-response curve fitted with a three-parameter Hill curve. The number in the top left corner of each plot is the R^2^, which measures the quality of the fitted curve. hr, hours; nL, nanoliter.

**Figure 5:**
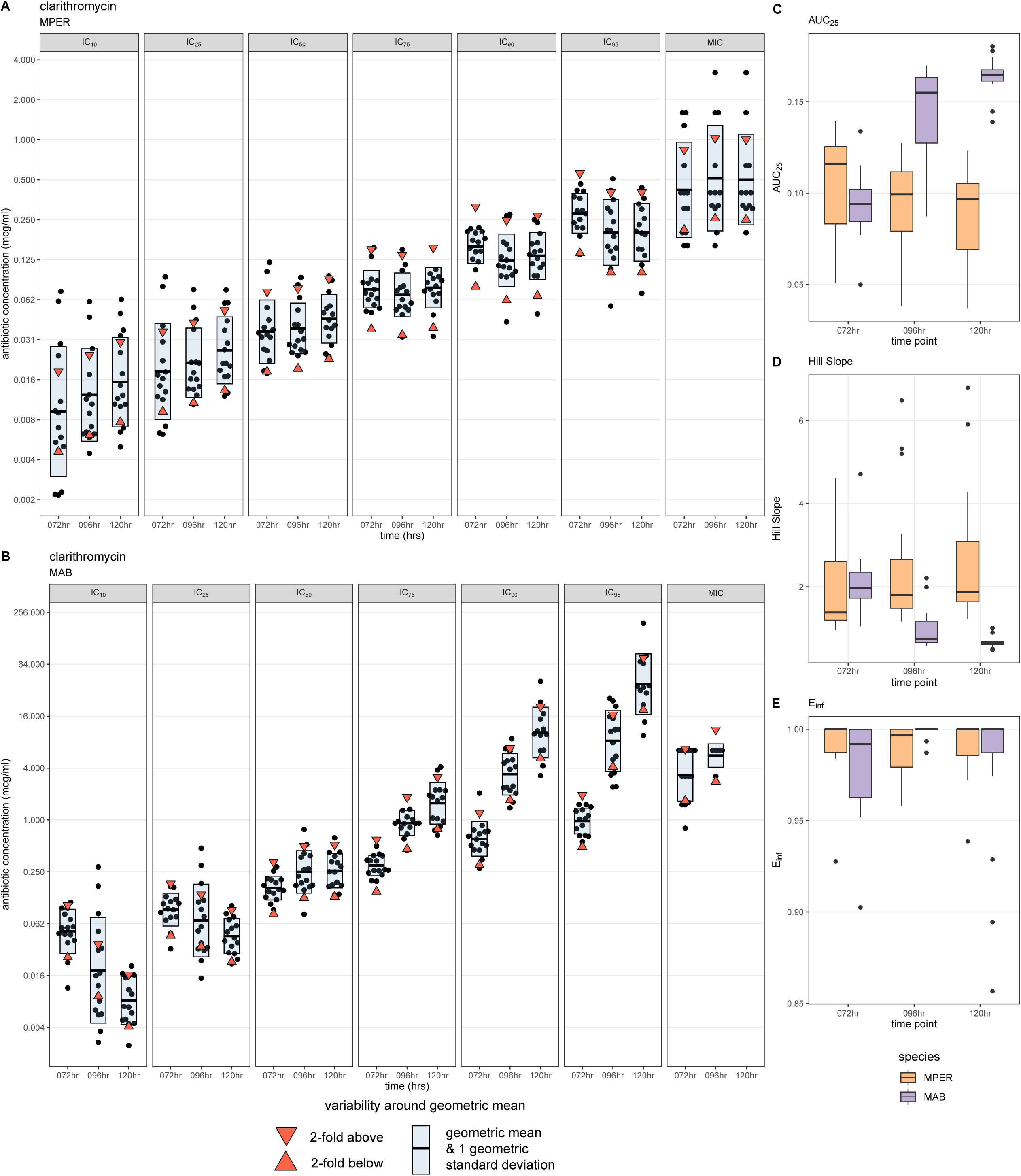
Dose-response metrics show inducible resistance to clarithromycin in *Mycobacterium abscessus*. The IC values for each replicate and the MICs for cefoxitin at each time point (black points) for *Mycobacterium peregrinum* (A) and *Mycobacterium abscessus* (B), with the geometric mean represented by the horizontal bar. The upper and lower limits of ±1 2-fold range around the geometric mean are represented by the downward- and upward-pointing red triangles, respectively. The blue shaded area is 1 geometric standard deviation around the geometric mean. Metrics from the dose-response curve can also show evidence of inducible resistance as seen with boxplots of the Area Under the Curve at 25% growth inhibition (AUC_25_, C) and the Hill Slope (D). These are not seen with the E_inf_ (maximum effect at infinite concentration). MPER, *Mycobacterium peregrinum*; MAB, *Mycobacterium abscessus* (B); hr, hours; IC_10,_ inhibitory concentration at 10% growth inhibition; IC_25_, inhibitory concentration at 25% growth inhibition; IC_50_, inhibitory concentration at 50% growth inhibition; IC_75_, inhibitory concentration at 75% growth inhibition; IC_90,_ inhibitory concentration at 90% growth inhibition; IC_95_, inhibitory concentration at 95% growth inhibition; MIC, minimum inhibitory concentration.

## Discussion

In this study, we demonstrate that antibiotic activity can be measured using dose-response curves, yielding metrics that are less variable than the MIC and that can identify inducible resistance at five days. AST that can provide precise strain-specific measurements of antibiotic activity have the potential to transform the treatment of drug-resistant infections through improved antibiotic selection and individualized antibiotic dosing.^22^ The consequences of flaws in MIC-based AST are best seen with MAB, where the absence of reliable and predictive metrics has hindered the search for effective and safe treatment for patients infected with this highly drug-resistant pathogen.^1^ We believe that AST performed using dose-response curves may have two potential advantages over MIC-based AST methods.

First, we show that the IC_50_ and IC_75_ allow precise measurement of antibiotic activity over an 8,000-fold range in concentration, in contrast to existing AST methods that approximate the MIC over a narrower range (8-500-fold for NTM rapid-grower AST panels). This precision has the potential to “personalize” antibiotic dosing and thereby benefit patients by lowering the risk of side effects while preserving antibiotic efficacy.^22,26^ In addition, increased accuracy of IC_50_ may lower the number of replicates needed when performing new AST validations, thereby allowing the faster incorporation of AST methods for new antibiotics into clinical microbiology laboratories.^39^

Second, we identify time-dependent changes in dose-response curve metrics that enabled the identification of inducible clarithromycin resistance with only five days of incubation. Existing MAB AST can identify inducible resistance to clarithromycin in only ∼60% of isolates after five days of incubation, necessitating the use of either extended incubation to fourteen days or molecular assays (*erm*(41) sequencing or line probe assays), which are usually only available at reference laboratories.^40^ We propose that an increase in the IC_75_, IC_90_, IC_95_, and AUC_25,_ and a decrease in the Hill slope between days three and five, could serve as a marker of inducible resistance and obviate the need for extended incubation or use of molecular methods to detect inducible clarithromycin resistance in MAB. This could allow the faster identification of inducible resistance and reduce delays in treatment initiation.

Our work has many strengths that make it clinically relevant. We used CLSI standards for antibiotic stock preparation and storage.^27^ We also performed experiments starting with an inoculum equivalent to 0.5 McFarland standard, used freshly prepared cefoxitin and CA-MHB with each experimental run, and incubated plates at 30°C for 3-5 days, all steps recommended by CLSI.^17,20^ We tested drugs that are recommended for use or appear promising for the treatment of MAB.^1,3,32,33,35^ R^2^ values served as a good measure of the quality of dose-response curves and might be incorporated into clinical microbiology quality control methods for dose-response curve AST. We were able to obtain results for omadacycline, which has been challenging with conventional AST due to trailing endpoints.^34,41^ We quantified the variability between different IC and MIC values, time points, and antibiotics. By incorporating dose-response curve metrics (AUC_25_ and the Hill slope), we compared antibiotic potency across species, drugs, and time points, a feature not currently possible with MICs or IC values.

Limitations of our work revolved around technical challenges. Drug-specific variability in IC and MIC values, as well as a low percentage of successful attempts, was likely due to one or more of a combination of antibiotic insolubility (clofazimine), the use of high antibiotic concentrations (SQ109), and low dispense volumes (bedaquiline and tigecycline), suggesting the need for optimization of antibiotic preparation for future experiments. Previous work has demonstrated the limited reproducibility of NTM AST results, even when testing is performed by reference laboratories, suggesting that some of the challenges we faced may be due to MAB itself.^42–44^ While we replaced the optically clear plate seal at each time point before obtaining OD_600_ values, condensation under the seal due to temperature differences between the incubator, biosafety cabinet, and ambient air may have introduced error in OD_600_ measurements.

The MIC and IC values we obtained were higher than expected, likely because the optical plate reader is more sensitive than the unaided eye in measuring bacterial growth, although errors in inoculum preparation cannot be ruled out. We excluded nearly 1/3 of MIC measurements because we could not obtain 100% growth inhibition, suggesting that new susceptibility breakpoints based on IC values will need to be established for each antibiotic-bacterial pair. Further validation of this work in benchmarked clinical isolates with different resistance mechanisms and a range of MIC values is needed. Antibiotic dosing with IC_50_ values and antibiotic selection with Hill metrics will need to be validated in animal and clinical studies.

In 1992, Soothill et al used *Pseudomonas aeruginosa* and gentamicin dissolved in solid agar plates at 12 different concentrations and demonstrated that the IC_50_ was significantly less variable than the MIC.^26^ Using CLSI conditions for rapid grower NTM AST, 384-well plates, and a digital drug dispenser, we redemonstrate Soothill et al’s findings more than thirty years later. Our results indicate that measuring antibiotic activity against MAB using dose-response curves can overcome the limitations of the MIC while allowing the rapid identification of inducible resistance to clarithromycin. These findings are applicable to other pathogens and have the potential to enable strain-specific dosing of antibiotics, facilitate the rapid detection of inducible resistance, and enhance antibiotic selection for the treatment of drug-resistant infections.

## Acknowledgments

HP was funded by 1K12TR004384 and 1K08AI190124. CLD serves on advisory boards for the following AN2, AstraZeneca, Galapagos, Grifols, GSK, Hyfe, Insmed, MannKind, Matinas BioPharma Holdings, Inc., Microbion, MicuRx, NobHill, Paratek Pharmaceuticals, Shionogi, Spero Therapeutics, Ostuka Pharmaceutical, Bill and Melinda Gates Foundation. He has received funding from AN2 Therapeutics, Bugworks, COPD Foundation, Cystic Fibrosis Foundation, Insmed, Juvabis, MannKind, Paratek Pharmaceuticals, Renovion, Spero, and Verona. AA received an Infectious Diseases Society of America G.E.R.M grant in 2024. HP would like to thank multiple people: Ms. Hanna Clutterbook-Cook for feedback on the manuscript draft, Dr. Robert Bonomo for the recommendation to use cefoxitin sodium rather than cefoxitin, Dr. Perry Riggle from Tufts Media Kitchen for preparing fresh media for each experimental run, and Mr. Hidetomi Nitta for creating a graphical user interface to review dose-response curves.

---

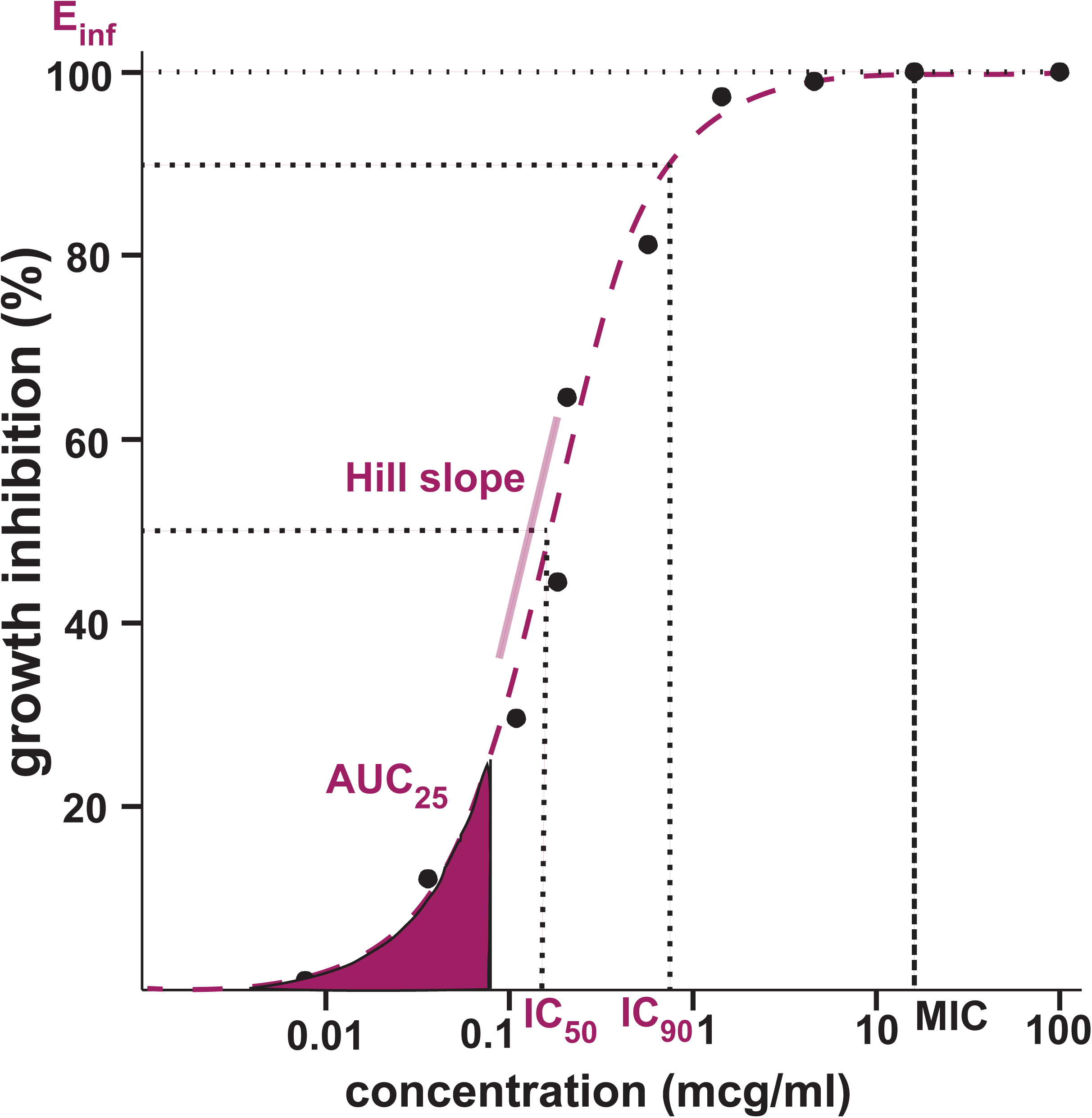

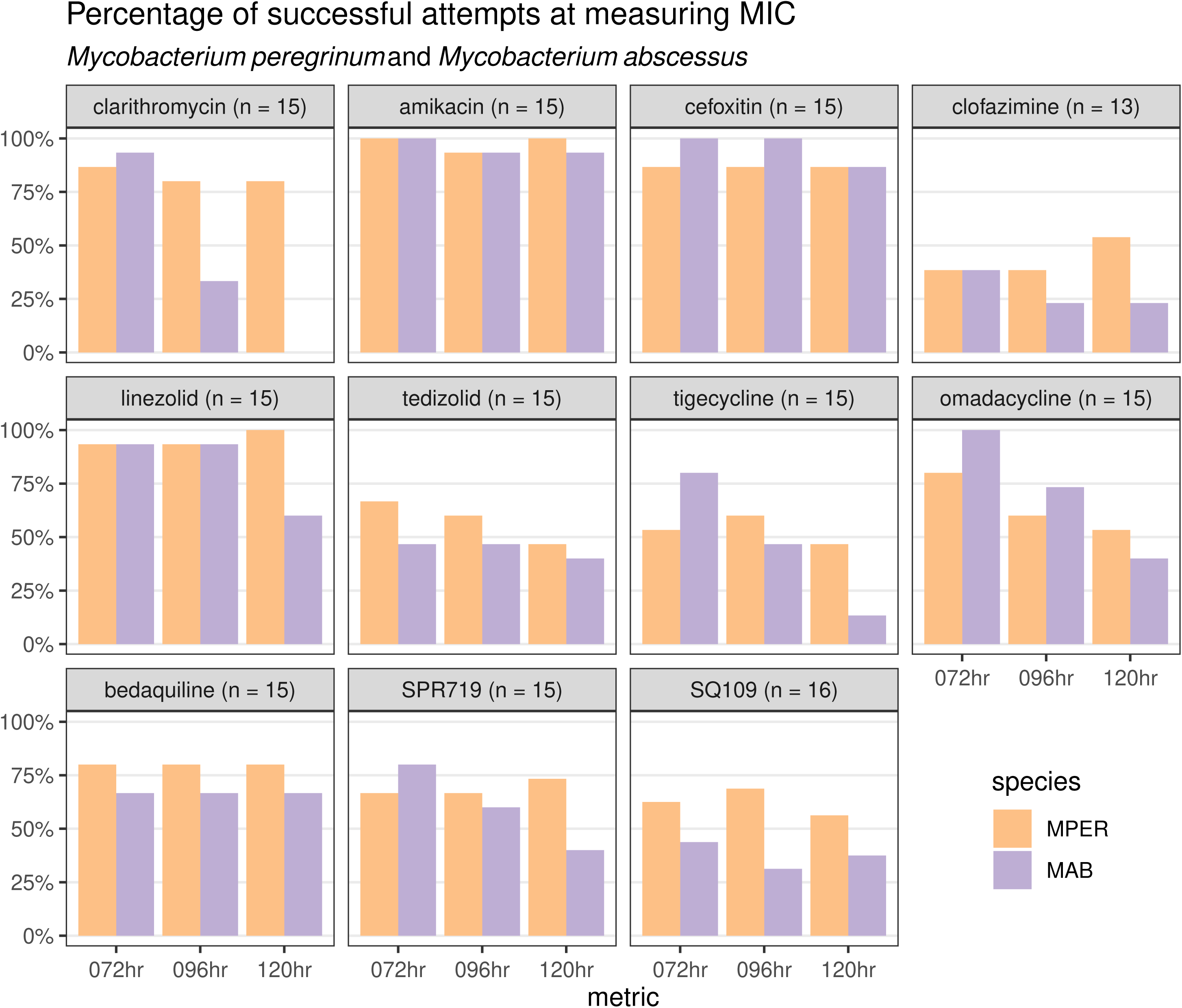

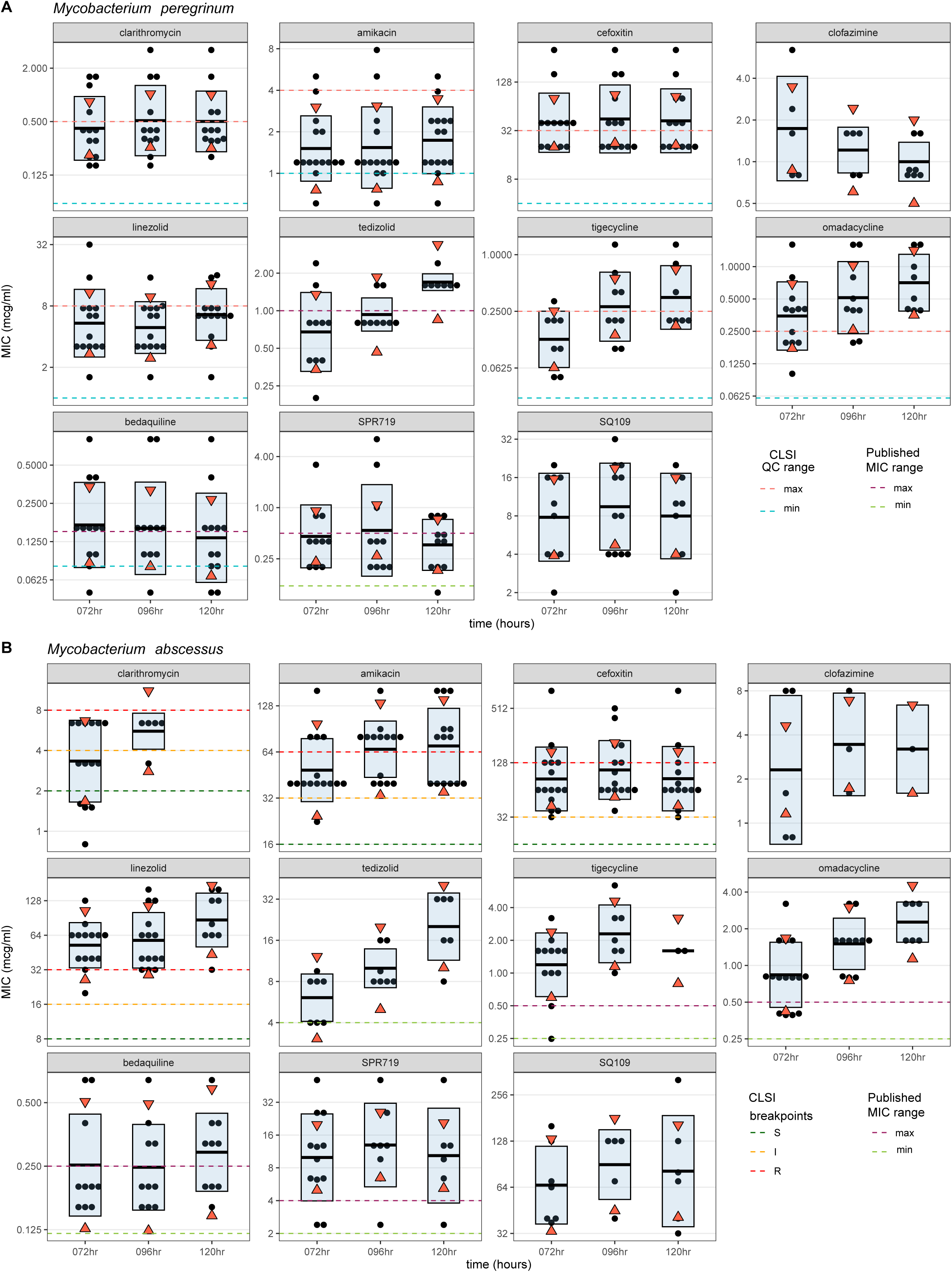
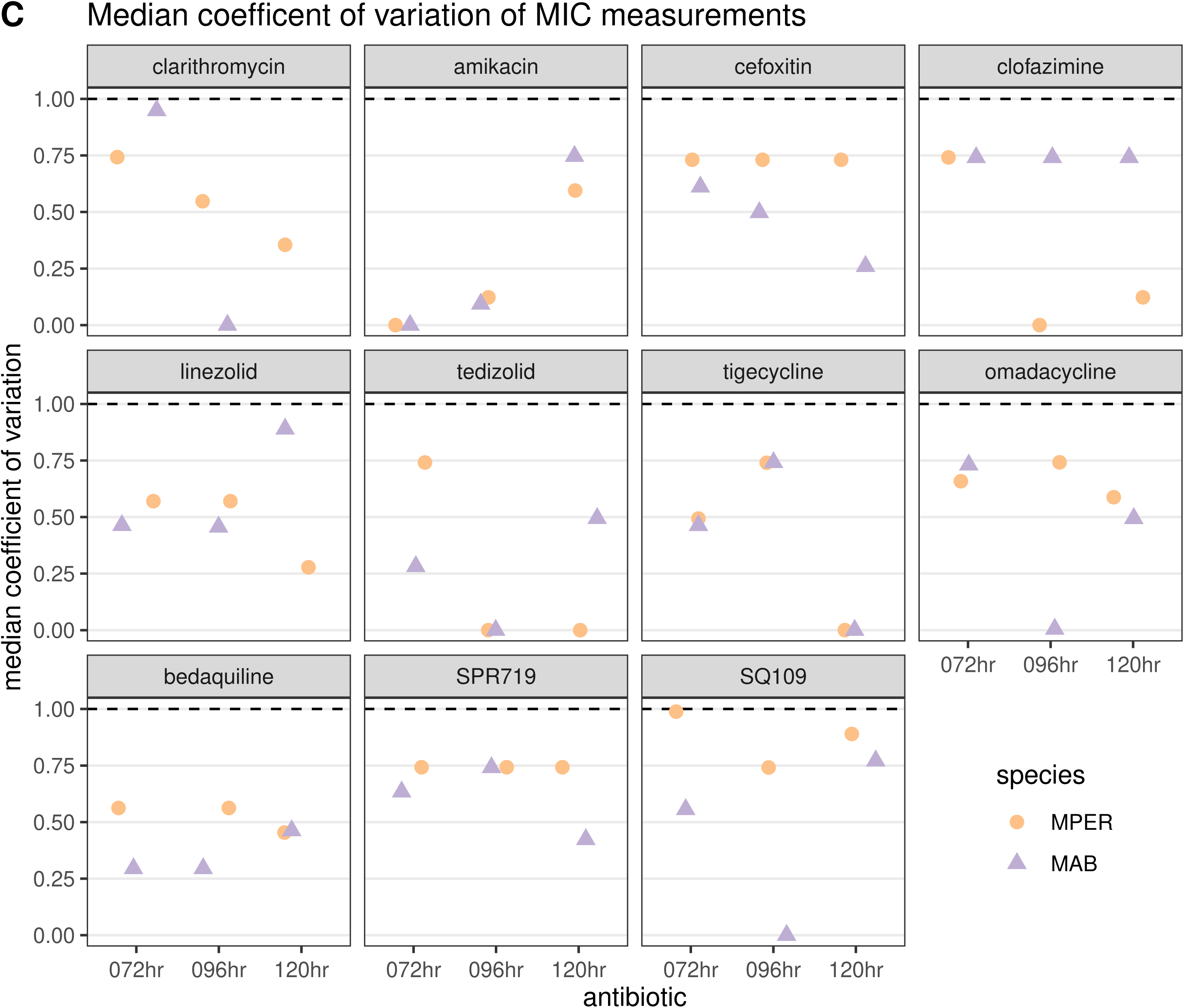

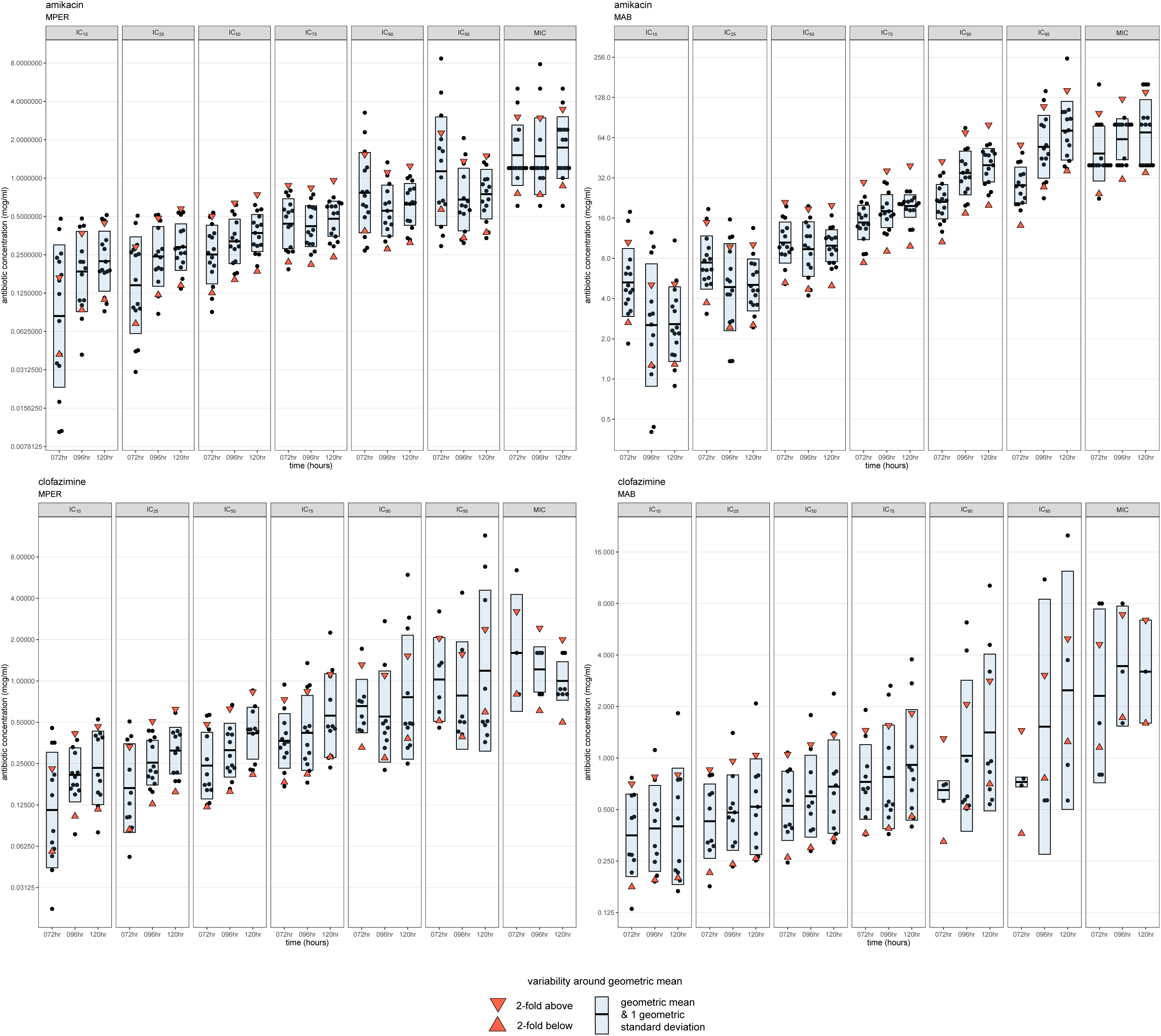

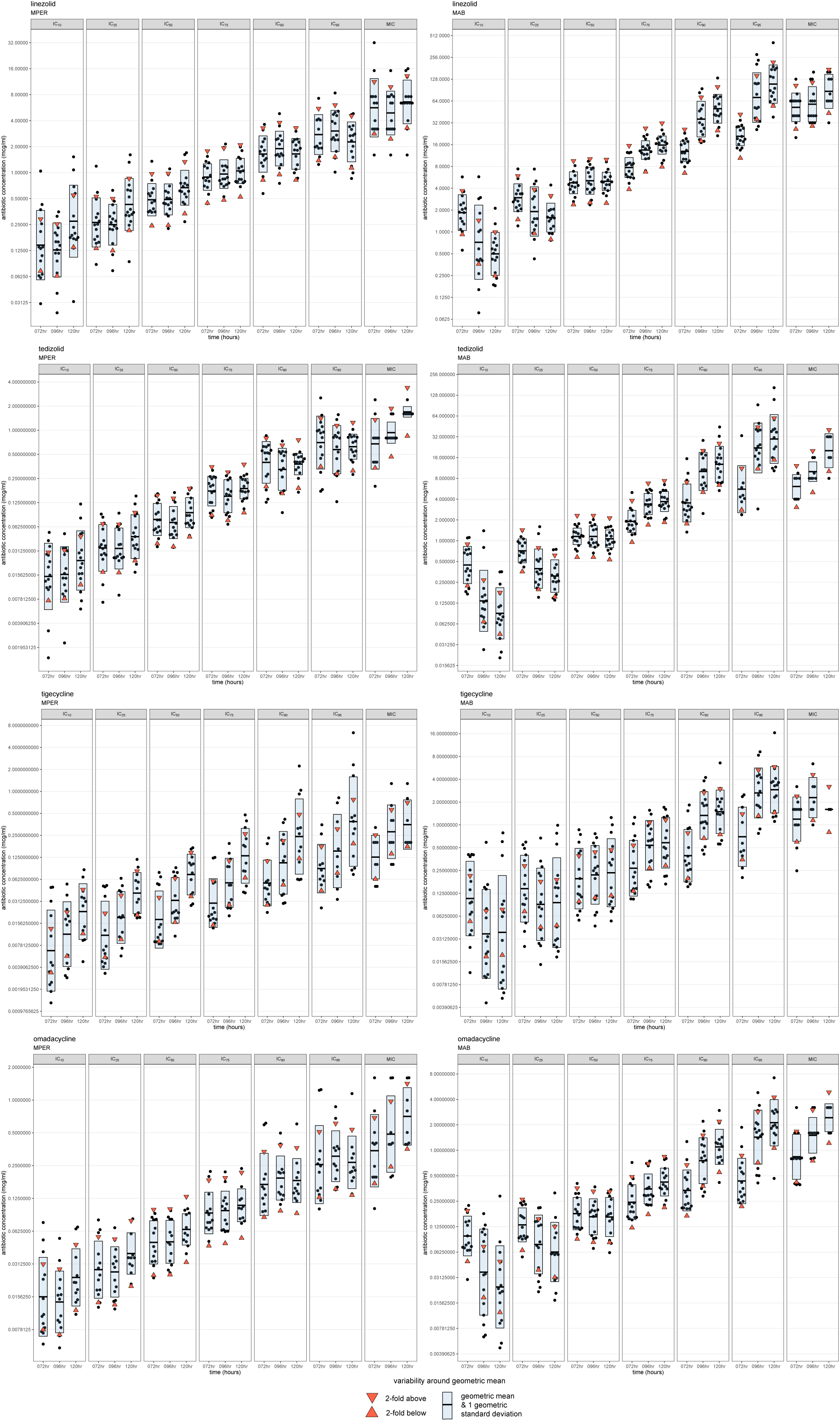

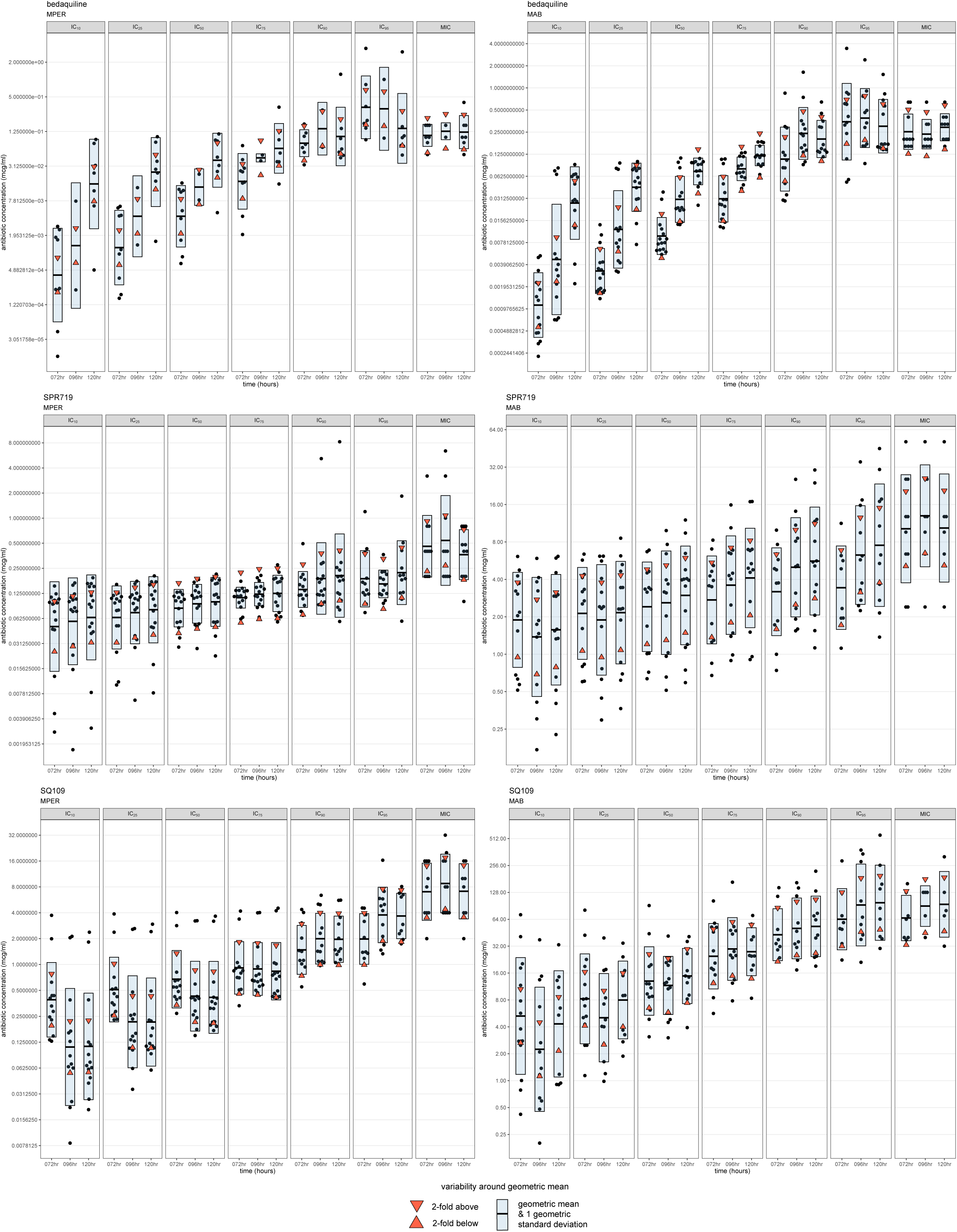

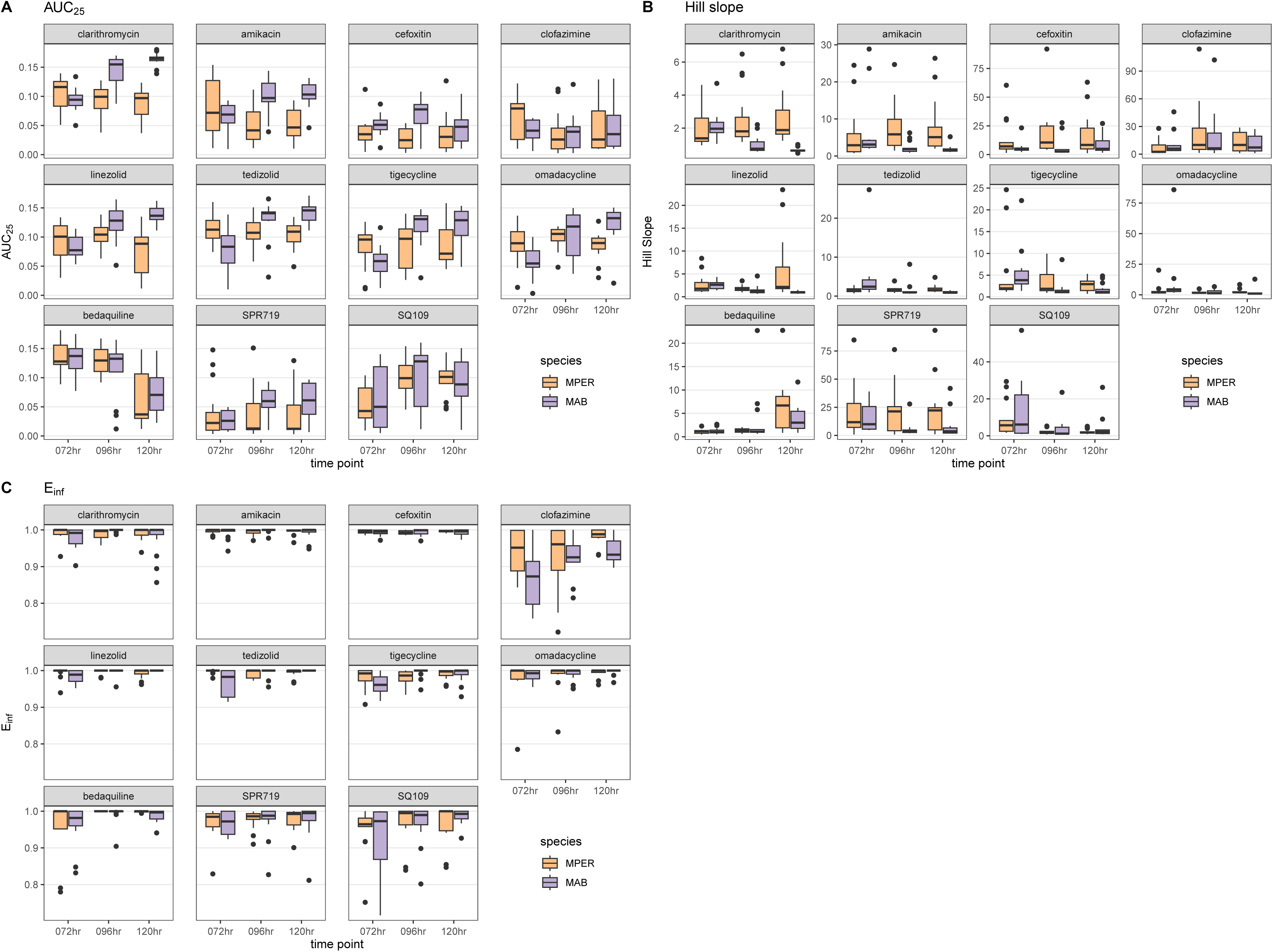

